# Monocyte-derived cells but not Microglia cause Oxidative Tissue Damage in Neuroinflammation

**DOI:** 10.1101/2024.09.18.612891

**Authors:** Juan Villar-Vesga, Donatella De Feo, Pauline Clément, Florian Ingelfinger, Can Ulutekin, Jeanne Kim, Maria Pena-Francesch, Katarina Wendy Schmidt, Elèni Meuffels, Viola Bugada, Deborah Greis, Sara Costa-Pereira, Laura Oberbichler, Frauke Seehusen, Francesco Prisco, Urvashi Dalvi, Christian Münz, Aiman S. Saab, Burkhard Becher, Sarah Mundt

## Abstract

Multiple sclerosis (MS) is characterized by neuroinflammation, oxidative stress, iron toxicity and mitochondrial dysfunction. Reactive oxygen species (ROS) produced by mononuclear phagocytes (MPs) are widely held to drive tissue damage, yet the specific roles of central nervous system (CNS)- resident versus CNS-invading MPs remain unclear. Here, by combining single-cell profiling with conditional gene targeting, we systematically dissected and interfered with ROS production across CNS MPs in a preclinical model for neuroinflammation. We show that CNS-invading monocyte derived cells (MdCs) exhibit a higher oxidative stress gene signature and produce more ROS compared to CNS-resident microglia. While NADPH oxidase 2 (NOX2), a phagocytic source of ROS, proved redundant, our findings underscore the critical role of mitochondrial ROS (mtROS) in driving oxidative tissue damage. Quenching mtROS through mitocatalase overexpression in MdCs, but not microglia, significantly alleviated neuroinflammation in mice. Thus, our study resolves a longstanding controversy, identifying MdCs as the primary driver of ROS-mediated neuropathology.

## Main

Multiple Sclerosis (MS) is the most common chronic inflammatory, demyelinating disorder of the central nervous system (CNS) affecting nearly 3 million people worldwide. The pathogenesis of MS is thought to be initiated by autoreactive T cells, while tissue damage is primarily driven by infiltrating and/or resident mononuclear phagocytes (MPs)^1,2^. Over time, accumulating pathological tissue damage often culminating in neurodegeneration and severe neurological disability^1^. Both studies in MS patients and preclinical models have consistently shown a strong correlation between oxidative stress and CNS tissue damage^3–6^.

Oxidative stress arises from an imbalance between the production of reactive oxygen species (ROS) by oxidant enzymes and the detoxifying effects of antioxidant enzymes^3,7^. Elevated ROS levels, including oxygen and nitrogen free radicals, oxidize lipids in cellular membranes, leading to their destabilization and damage^7,8^. Furthermore, protein nitration, particularly on tyrosine residues, can alter protein function^9^. The importance of oxidative tissue damage in MS is highlighted by the oxidation of lipids^5^ and nucleic acids^4^, as well as protein nitration observed in MS patient brain lesions^8^. Efforts to reduce ROS levels have been shown to alleviate disease severity in preclinical MS models^3,10,11^, suggesting targeting ROS production or enhancing antioxidant activity as a promising therapeutic approach in MS^7^.

CNS-resident microglia and border-associated macrophages (BAMs) as well as infiltrating monocyte-derived cells (MdCs) have been implicated in ROS production and oxidative injury during CNS neuroinflammation^3,4,12–16^. However, the specific contribution of these distinct cellular subsets to ROS mediated tissue damage remains unclear. Recent studies have highlighted the role of mitochondrial complex I (mtCI) activity in driving ROS production by MPs during neuroinflammation^6^, yet the cell-specific contributions of resident versus invading phagocytes were not addressed.

To address this gap, we employed single-cell profiling to characterize phagocytic ROS production in MPS during human and murine CNS neuroinflammation. In combination with conditional gene targeting in mice, we identified MdCs as the primary drivers of ROS mediated tissue damage in CNS neuroinflammation.

## Results

### Human and Murine Neuroinflammation is associated with a high ROS signature in Damage-Associated MPs

First, to map oxidative stress at the single-cell level, we integrated the immune compartments of five publicly available human single nuclei RNA sequencing (snRNA-seq) MS datasets^17–21^ and interrogated the combined data for the expression of genes associated with oxidative stress. Therefore, we computed a “ROS Enrichment Score” (human GO:0072593 reactive oxygen species metabolic process) within the MP clusters (Fig. 1a). Briefly, cells were subsetted based on immune-related annotations from the original datasets. Immune cells were integrated using Harmony to mitigate batch effects, and clustered using the Louvain algorithm. A curated list of established marker genes for each cell type was used to manually annotate the clusters. UMAP was employed to visualize clusters and cell type annotations across control and MS samples (Extended Data Fig. 1a-c,e). For each cell annotated as an MP, the “ROS Enrichment Score” was computed. We found an increase in the average module score of MPs from MS patients compared to controls (Fig. 1b, Extended Data Fig. 2d) particularly associated with areas containing white matter lesions (Lesion Edge (LE) and Active Lesion) (Fig. 1c).

**Figure 1.**
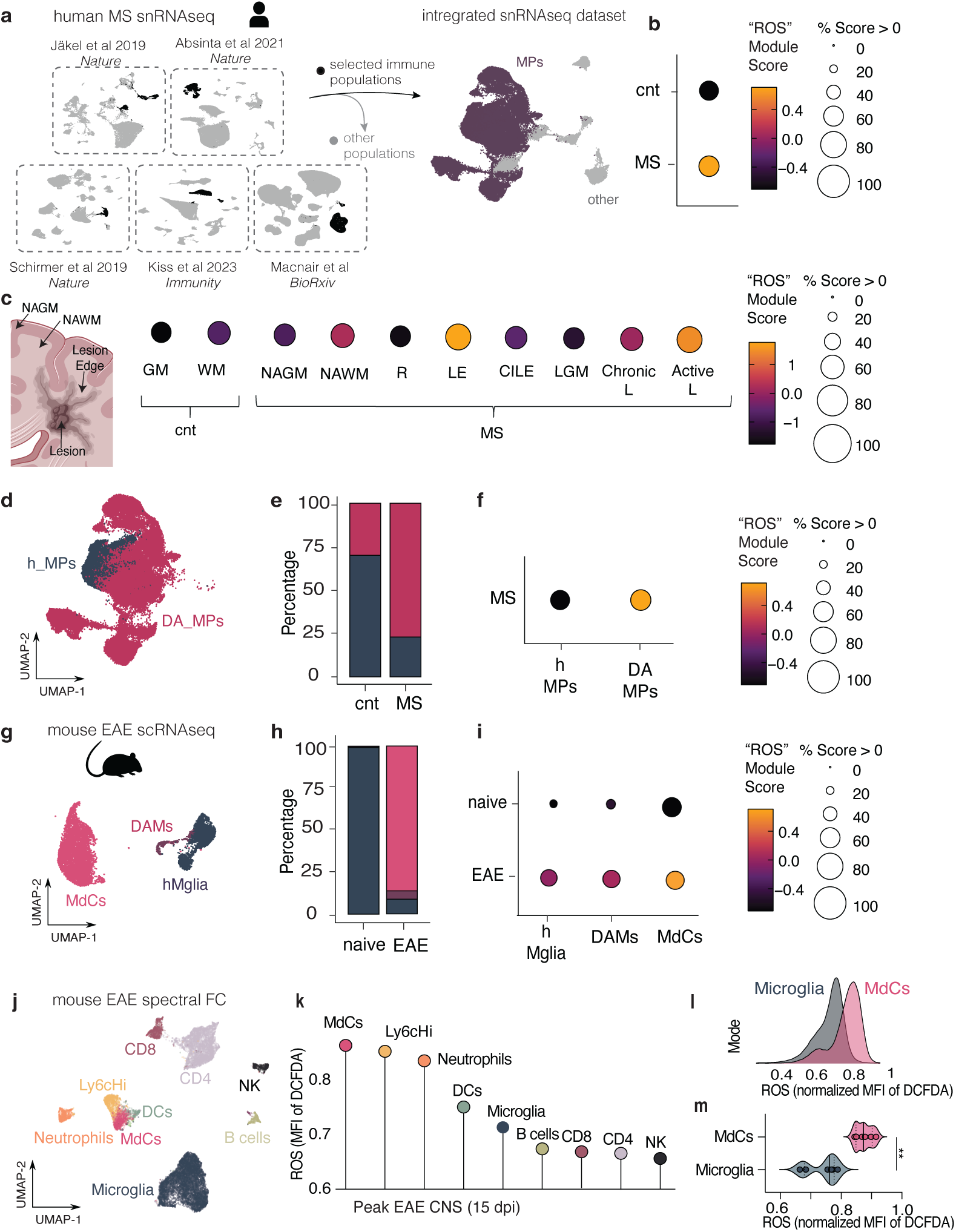
Human and murine neuroinflammation is associated with a high ROS signature in damage-associated MPs. **a,** Schematic illustration of cluster selection and data integration of multiple human snRNA-seq datasets (author and date depicted) **b,** Dot plot illustrating *“ROS”* module score in mononuclear phagocytes (MPs) from control and MS patient samples. **c,** Dot plot of the *“ROS”* module score values in MPs across different tissues from control and MS patients. Control tissues: GM (Gray Matter), WM (White Matter); MS tissues: NAGM (Normal Appearing GM), NAWM (Normal Appearing WM); R (Remyelination), LE (Lesion Edge), CILE (Chronic Inactive LE), Lesion GM, Chronic Lesion, Active Lesion. **d,** UMAP of the two clusters of MPs: damage-associated MPs (DA-MPs) and homeostatic-associated MPs. **e,** Bar graph of the percentage of the MP clusters in control and MS samples. **f,** Dot plot illustrating *“ROS”* module score values in the two MPs clusters. **g,** UMAP of immune clusters found in Mendiola *et al*. **h,** Bar graph of the percentage of the immune clusters found in control and EAE CNS. **i,** Dot plot illustrating *“ROS”* module score (mouse GO term: 0072593) in homeostatic microglia (hMicro), damage-associated microglia (DAMs), and monocyte-derived cell clusters (MdCs). **k,** UMAP of the CNS immune cell populations found during peak EAE **k,** Lollipop plot of DCFDA expression across different immune clusters in the inflamed CNS (peak EAE CNS day 15 post-immunization). **l,** Histogram showing DCFDA expression in MdCs and microglia. **m,** Violin plot of DCFDA expression in MdCs and microglia (control n=6, EAE n=6). (unpaired two-tailed t-test; **P = 0.0071). Pooled data from three experiment. For **b,c,f,i,** Circle size is depicting % of normalized expression detected per cluster and color is depicting expression level as Average Module Score or Average Expression.

Additional sub-clustering of the MPs revealed cluster families with higher scores for a homeostatic gene signature (H-MPs, clusters 1 and 13) and a damage-associated signature (Fig. 1d, Extended Data Fig.1e). The latter family comprised damage associated microglia (DAM) and MdC genes (Extended Data Fig.1b-d; DAM genes: *APOE, LPL, TREM2, PLIN3, C1QA, C1QC* and *MdC: CD14, FCGR3A, CD36, FCGR1A, ITGAM, ITGAX*) and consequently showed a higher abundance in MS patients compared to controls (Fig. 1d-e). The “ROS Enrichment Score” was clearly increased in the DA-MPs (Fig. 1f) suggesting that ROS production is elevated in MPs during human neuroinflammation.

To investigate the oxidative stress response in MPs during experimental neuroinflammation, we re-interrogated three publicly available scRNA-seq data of CNS immune cells of mice with experimental autoimmune encephalomyelitis (EAE)^3,6,22^. Briefly, for each dataset, CNS immune cells were individually re-clustered using the Louvain algorithm and manually re-annotated with the same marker genes to ensure consistency across all datasets. The resulting clusters and cell type annotations were visualized in UMAPs. In contrast to human data, a clear separation between tissue resident and infiltrating phagocytes can be accomplished in the murine setting. Three MP clusters were annotated: homeostatic microglia (hMicroglia), DAMs, and MdCs (Fig. 1g, Extended Data Fig. 1f-g), the latter two clusters being enriched in the inflamed CNS (Fig. 1h). Computing the “ROS Enrichment Score” (mouse GO:0072593 reactive oxygen species metabolic process) confirmed a higher expression in MdCs over homeostatic microglia and even DAMs at peak EAE (Fig. 1i). In contrast to other MPs, the ROS signature in MdCs increased over time with highest expression in peak (Jordão *et al*. 2019^22^, GEO accession code GSE118948) (Extended Data Fig. 1h-i,l) and chronic phases of EAE (Peruzzotti-Jametti *et al.* 2024^6^, GEO:GSE248175) (Extended Data Fig. 1k,m).

We next asked whether the higher expression of transcripts related to the oxidative stress response translated into increased production of ROS *in vivo*. We immunized C57BL/6 mice with MOG_35-55_ peptide in Complete Freund’s Adjuvant (CFA) and pertussis toxin (PT) to induce active EAE. At peak EAE (day 15 post-immunization), we performed multiplexed flow cytometry (FC). Purified CNS leukocytes were stained with fluorchrome-tagged antibodies to identify major immune cell subsets in combination with dichlorodihydrofluorescein diacetate (DCFDA) to detect cellular oxidative stress^3^. We used FlowSOM to cluster distinct immune cell populations which were then projected onto a UMAP for dimensionality reduction (Fig. 1j, Extended Data Fig 1o). We determined active ROS production in individual clusters by analyzing the normalized mean fluorescent intensity of DCF. MdCs, Ly6C^high^ monocytes, and neutrophils were among the immune cell populations with the highest ROS intensity, while DCs, microglia and lymphocytes expressed lower levels of ROS (Fig. 1k, Extended Data Fig 1p). Direct comparison of FC raw data revealed a >5 fold higher ROS production in MdCs compared to microglia (Extended Data Fig. 1r). Together, these results show high oxidative stress signature and ROS production in MPs, specifically in CNS invading MdCs during neuroinflammation.

### *Cybb* gene deletion in CNS MPs does not affect neuroinflammation

Previous studies have suggested that phagocytes primarily produce ROS via the oxidative enzyme NADPH oxidase 2 (*NOX2/Cybb*)^3,12^. Consistent with this concept, *Cybb* deficient mice displayed ameliorated clinical scores in EAE^13^. Our analysis of murine scRNA-seq data revealed upregulated *Cybb* expression in both CNS-DAMs and MdCs during peak and chronic EAE (Extended Data Fig. 2a). To validate these findings, we FACS-sorted CNS phagocytes and confirmed that MdCs exhibited approximately 2.5-fold higher *Cybb* protein expression compared to microglia (Extended Data Fig. 2b).

To assess whether NOX2 plays a functional role on neuroinflammation, we conditionally ablated *Cybb*^15^ in MPs. Given the elevated oxidative stress gene signature, increased ROS levels, and higher *Cybb* expression in MdCs, we first utilized the inducible *Ccr2^CreERT2^* strain to specifically target monocytes and their progeny^23^. We induced active EAE in *Ccr2^CreERT2/+^ Cybb^fl/fl^* and *Ccr2^+/+^ Cybb^fl/fl^* littermate control mice (Fig. 2a). To prevent early interference with NOX2 expression, which could influence the priming of autoreactive T cells^24^, we administered tamoxifen after disease onset to induce Cre-mediated recombination. We confirmed the specific deletion of *Cybb* in MdCs by qPCR of isolated cell populations (Fig. 2b). However, no differences in EAE disease severity or CNS immune cell infiltration (CD45+ CD44+ cells, see gating strategy in Extended Data Fig. 1q) were observed between *Ccr2^CreERT2/+^ Cybb^fl/fl^* and *Ccr2^+/+^ Cybb^fl/fl^* littermate controls (Fig. 2c-e).

**Figure 2.**
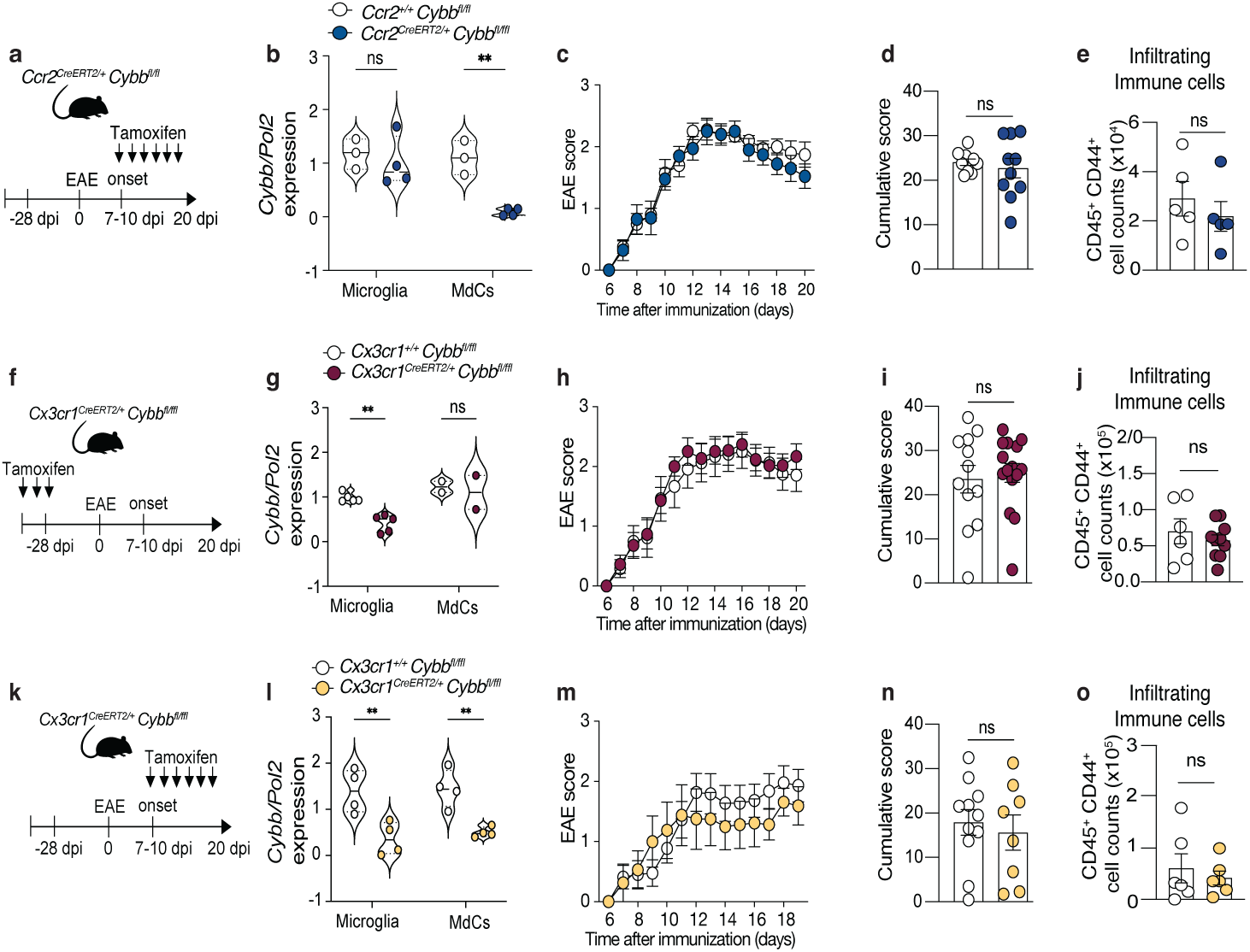
NOX2 plays a redundant role in MP mediated oxidative tissue damage during neuroinflammation: **a,** Schematic illustration of the tamoxifen treatment strategy to target *Cybb* in MdCs during neuroinflammation. **b**, Relative gene expression of *Cybb*/*Pol2* in CNS microglia and MdCs in *Ccr2^+/+^ Cybb^fl^*^/fl^ (n=3, f) and *Ccr2 ^CreERT2/+^ Cybb^fl^*^/fl^ (n=4, f) mice (unpaired two-tailed t-test, **p=0.0043 in MdCs). **c-d**, Active EAE was induced by MOG_35-55_/CFA/PT immunization. Plots depict mean clinical score (**c**) over time (two-way ANOVA, and Sidak’s post hoc test) and (**d**) cumulative clinical score *Ccr2^+/+^ Cybb^fl^*^/fl^ (n=11, m/f) and *Ccr2 ^CreERT2/+^ Cybb^fl^*^/fl^ (n=10, m/f) mice (unpaired two-tailed t-test). **e,** Absolute counts of CNS infiltrating immune cells (gated in FlowJo CD45+CD44+, gating strategy Extended Data Fig. 1q) at 19 d.p.i. in *Cx3cr1^+/+^ Cybb^fl^*^/fl^ (n=5, f) and *Cx3cr1^CreERT2/+^ Cybb^fl^*^/fl^ (n=4,f) mice. Unpaired two-tailed t-test. **f,** Schematic representation of tamoxifen treatment to target embryonic hematopoiesis-derived macrophages, but not bone marrow derived MdCs. **g,** Relative gene expression of *Cybb*/*Pol2* in CNS microglia and MdCs in *Cx3cr1^+/+^ Cybb^fl^*^/fl^ (Microglia n=6, MdCs=2 f) and *Cx3cr1 ^CreERT2/+^ Cybb^fl^*^/fl^ (Microglia n=5, MdCs n=2, f) mice (unpaired two-tailed t-test, **p=0.0043 in Microglia). **h-i,** Active EAE was induced by MOG_35-55_/CFA/PT immunization. Plots show mean clinical score over time (Two-way ANOVA, and Sidak’s post hoc test) (**h**) and cumulative clinical score (**i**) in *Cx3cr1^+/+^* mCAT^fl/wt^ (n=15, m/f) and *Cx3cr1^CreERT2/+^* mCAT^fl/wt^ (n=15, m/f) mice (unpaired two-tailed t-test). **j,** Absolute counts of CNS infiltrating immune cells (gated in flowjo CD45+CD44+, gating strategy Extended Data Fig. 1q) at 19 d.p.i. in *Cx3cr1^+/+^ Cybb^fl^*^/fl^ (n=6, f) and *Cx3cr1 ^CreERT2/+^ Cybb^fl^*^/fl^ (n=10, f) mice. Unpaired two-tailed t-test. **k**, Schematic illustration of the tamoxifen treatment strategy to target *Cybb* in microglia and MdCs during neuroinflammation. **l**, Relative gene expression of *Cybb*/*Pol2* in CNS microglia and MdCs in *Cx3cr1^+/+^ Cybb^fl^*^/fl^ (n=4, f) and *Cx3cr1 ^CreERT2/+^ Cybb^fl^*^/fl^ (n=4, f) mice (unpaired two-tailed t-test, **p=0.0036 in Microglia and **p=0.0063 in MdCs). **m-n**, Active EAE was induced by MOG_35-55_/CFA/PT immunization. Plots depict mean clinical score (**m**) over time (two-way ANOVA, and Sidak’s post hoc test) and (**n)** cumulative clinical score *Cx3cr1^+/+^ Cybb^fl^*^/fl^ (n=11, m/f) and *Cx3cr1 ^CreERT2/+^ Cybb^fl^*^/fl^ (n=8, m/f) mice (unpaired two-tailed t-test). **o,** Absolute counts of CNS infiltrating immune cells (gated in FlowJo CD45+CD44+, gating strategy Extended Data Fig. 1q) at 19 d.p.i. in *Cx3cr1^+/+^ Cybb^fl^*^/fl^ (n=6, f) and *Cx3cr1 ^CreERT2/+^ Cybb^fl^*^/fl^ (n=6, f) mice.

It is important to note that the active EAE mouse model relies on the subcutaneous deposition of MOG/CFA which may lead to continuous T cell activation in the periphery. We speculated that this ongoing influx of encephalitogenic T cells could overshadow any subtle effects of ROS depletion on clinical outcomes. To address this, we performed an adoptive transfer of encephalitogenic T cells to induce passive EAE in *Ccr2^CreERT2/+^ Cybb^fl/fl^* and littermate control mice. Despite this approach, we observed no impact of *Cybb* deletion on disease progression in this model either (Extended Data Fig. 2c-d).

While our data suggest a predominant role of MdCs in regulating and producing ROS (Fig. 1i-k), previous studies have highlighted a key role for microglia in ROS production during CNS neuroinflammation^3,6,12–15,25^. To evaluate the impact of NOX2 mediated ROS production in microglia, we crossed the *Cybb^fl^* allele to the *Cx3cr1^CreERT2^* strain^26^ (Fig. 2f). To induce *Cybb* ablation specifically in microglia (and BAMs), we treated *Cx3cr1^CreERT2/+^ Cybb^fl/fl^* mice with tamoxifen (Fig. 2f, arrows) 4-5 weeks before disease induction allowing time for bone marrow-derived cells to replace MdCs^26^. Tamoxifen treated *Cx3cr1^+/+^ Cybb^fl/fl^* mice served as the control group. After confirming successful *Cybb* depletion in microglia (Fig. 2g), we induced active EAE through MOG/CFA/PT immunization. However, the deletion of *Cybb* in microglia had no effect on clinical disease severity, as *Cx3cr1^CreERT2/+^ Cybb^fl/fl^* mice exhibited similar cumulative disease scores and immune cell infiltration compared to *Cx3cr1^+/+^ Cybb^fl/fl^* controls (Fig. 2h-j).

To assess whether MdCs and microglia play a redundant role in ROS-mediated tissue damage, we simultaneously ablated *Cybb* in both populations by continuously administrating tamoxifen to *Cx3cr1^CreERT2/+^ Cybb^fl/fl^* mice upon disease onset (Fig. 2k). While we confirmed a significant reduction of *Cybb* expression in both microglia and MdCs (Fig. 2l), this approach also failed to affect clinical severity or CNS inflammatory cell infiltration (Fig. 2m-o).

To investigate why *Cybb* depletion did not alleviate neuroinflammation as previously reported^13,15^,we assessed its impact on ROS production using flow cytometry. Surprisingly, despite effective gene targeting of *Cybb*, we observed no change in cellular ROS levels (DCF expression) in either of our conditional models (Extended Data Fig. 2e-g). This suggests that while NOX2 has been described as a major oxidative enzyme in phagocytes, its loss may be compensated by alternative mechanisms of ROS production. In summary, *Cybb* depletion was insufficient to reduce ROS production in MPs, and thus, did not influence the disease outcome in EAE.

### Microglia play a redundant role in mtROS-mediated tissue damage

In addition to *Cybb* and other oxidant enzymes, recent studies have identified reverse electron transport (RET) as a key mechanism driving mitochondrial and general ROS upregulation during neuroinflammation^6^. RET is characterized by increased mitochondrial complex I (mtCI) activity, which catalyzes the formation of superoxide and peroxide. Reanalyzing the mtCI module score from a published scRNA-seq dataset^6^ showed high expression not only in DAMs but also specifically in MdCs during peak and in chronic stages of EAE (Extended Data Fig. 3a-b) suggesting this pathway to be active in both phagocyte subsets.

We next aimed to functionally target mtROS in CNS MPs during neuroinflammation. To achieve this, we employed the genetic overexpression of mitochondrial-tagged catalase (mCAT), which has previously been used to enhance antioxidant capacity^27,28^ and reduce mtROS production. To induce mCAT expression in MPs, we crossed the *Cx3cr1^CreERT2^* strain to the mCAT allele, which includes a floxed STOP cassette between the mCAT and the CAG promoter, as well as a human influenza haemagglutinin (HA) tag for detection via HA-specific antibodies. We induced active EAE in *Cx3cr1^CreERT^*^/+^ mCAT^fl/+^ mice and continuously treated them with tamoxifen upon disease onset to induce mCAT expression in CNS MPs as described above (Fig. 3a). Immunoblot analysis of cell lysates from FACS-purified MPs populations confirmed mCAT overexpression (Extended Data Fig. 3c). As expected, mCAT overexpression significantly reduced both cellular ROS and mtROS production in microglia and MdCs, as measured by DCFDA and mitosox-based flow cytometry analysis (Fig. 3b-c). This reduction in mtROS in CNS MPs was accompanied by an amelioration of neuropathology, as evidenced by lower cumulative disease scores (Fig. 3d-e) and reduced CNS immune cell infiltration (Fig. 3f).

**Figure 3.**
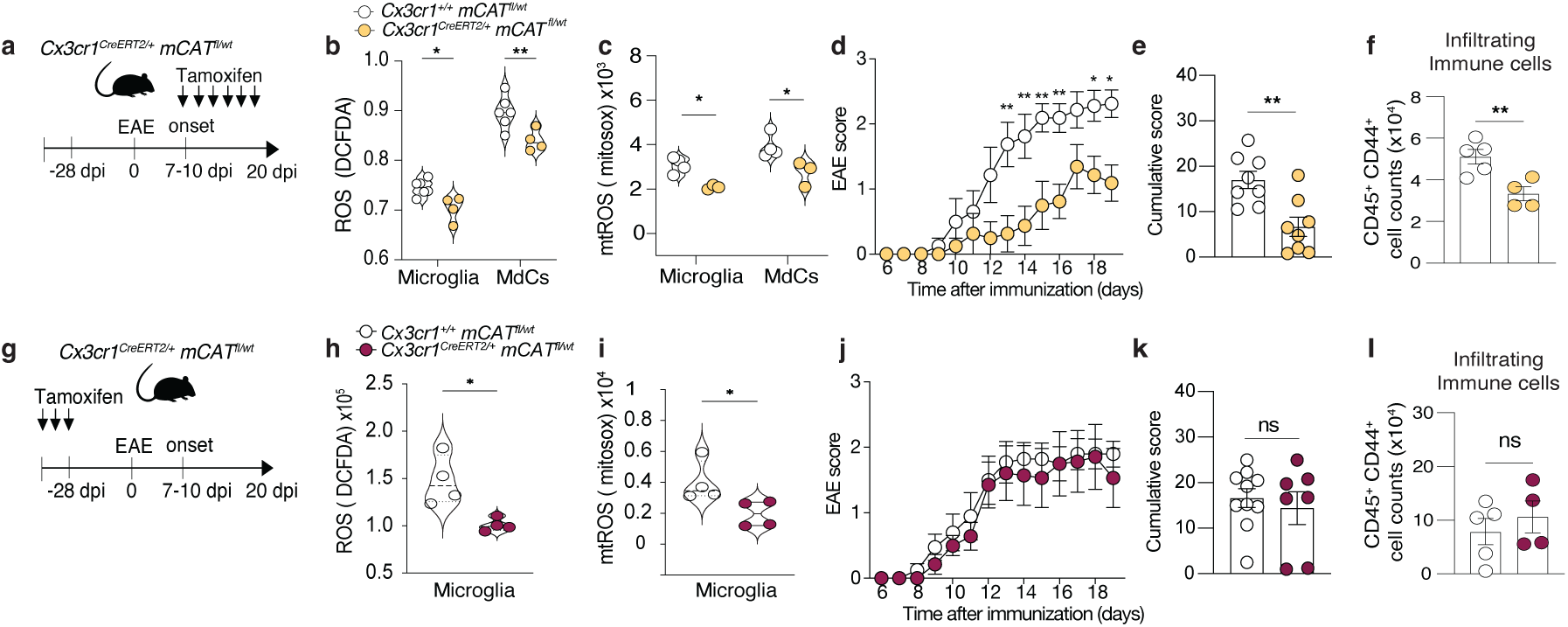
Microglia play a redundant role in mtROS mediated tissue damage. **a,** Schematic illustration of the tamoxifen treatment strategy employed to reduce mtROS in CX3CR1 expressing MPs. **b**, ROS production measured by normalized DCFDA expression in microglia and MdC clusters (generated in R) of *Cx3cr1^+/+^*mCAT^fl/wt^ (n=6, f) and *Cx3cr1^CreERT2/+^* mCAT^fl/wt^ (n=4, f). Pooled data from two experiments (unpaired two-tailed t-test, *P=0.04 in Microglia and **P=0.0071 in MdCs). **c**, Mitochondrial ROS (mtROS) production measured by mitosox expression in manually gated (FlowJo) microglia and MdC of *Cx3cr1^+/+^* mCAT^fl/wt^ (n=4, f) and *Cx3cr1^CreERT2/+^*mCAT^fl/wt^ (n=3 f). Unpaired two-tailed t-test (*P=0.0237 in Microglia and *P=0.0363 in MdCs). **d**-**e**, Active EAE was induced by MOG_35-55_/CFA/PT immunization. (**d**) Plots show mean clinical score over time (two-way ANOVA, and Sidak’s post hoc test, day 13 **P=0.0022, day 14 **P=0.0023, day 15 **P=0.0031, day 16 **P=0.0059, day 18 *P=0.0453, day 19 *P=0.0109) and (**e**) cumulative clinical score in *Cx3cr1^+/+^* mCAT^fl/wt^ (n=8, f) and *Cx3cr1^CreERT2/+^* mCAT^fl/wt^ (n=8, f) mice. Unpaired two-tailed t-test (*P=0.0405). **f,** Absolute counts of CNS infiltrating immune cells (gated in FlowJo CD45+CD44+, gating strategy Extended Data Fig. 1q) at 19 d.p.i. in *Cx3cr1^+/+^* mCAT^fl/wt^ (n=5, f) and *Cx3cr1^CreERT2/+^* mCAT^fl/wt^ (n=4, f) mice. Unpaired two-tailed t-test (*P=0.0091). **g,** Schematic representation of tamoxifen treatment to target embryonic hematopoiesis-derived macrophages, but not bone marrow derived MdCs**. h**, ROS production measured by DCFDA median expression in manually gated (FlowJo) microglia from *Cx3cr1^+/+^*mCAT^fl/wt^ (n=4, f) and *Cx3cr1^CreERT2/+^* mCAT^fl/wt^ (n=4, f)) in steady state CNS (unpaired two-tailed t-test; *P=0.0135). **i**, mtROS production measured as mitosox median expression in manually gated (FlowJo) microglia from *Cx3cr1^+/+^* mCAT^fl/wt^ (n=4, f) and *Cx3cr1^CreERT2/+^*mCAT^fl/wt^ (n=4, f) in steady state CNS (unpaired two-tailed t-test; *P=0.0423).**j-k,** Active EAE was induced by MOG_35-55_/CFA/PT immunization. Plots show mean clinical score over time (Two-way ANOVA, and Sidak’s post hoc test) (**j**) and cumulative clinical score (**k**) in *Cx3cr1^+/+^* mCAT^fl/wt^ (n=10, m/f) and *Cx3cr1^CreERT2/+^* mCAT^fl/wt^ (n=7, m/f) mice. Unpaired two-tailed t-test. Pooled data from two experiments. **l**, Absolute counts of CNS infiltrating immune cells (gated in FlowJo CD45+CD44+, gating strategy Extended Data Fig. 1q) at 19 d.p.i. in *Cx3cr1^+/+^*mCAT^fl/wt^ (n=5, f) and *Cx3cr1^CreERT2/+^* mCAT^fl/wt^ (n=4, f) mice. Unpaired two-tailed t-test.

To determine whether microglia are specifically responsible for the mtROS mediated tissue damage, we employed the early tamoxifen treatment in *Cx3cr1^CreERT^*^/+^ mCAT^fl/+^ inducing mCAT expression 4-5 weeks prior to disease onset (Fig 3g). We confirmed microglia-specific mCAT overexpression (Extended Data Fig. 3d), which led to a reduction in cellular ROS and mtROS before EAE induction (Fig. 3h-i). However, following MOG/CFA/PT immunization, there were no differences in disease severity or CNS immune cell infiltration compared to controls (Fig. 3j-l). We also evaluated myelin integrity in spinal cord sections but found no differences in demyelination between *Cx3cr1^CreERT^*^/+^ mCAT^fl/+^ and littermate controls (Extended Data Fig. 3e-f). Together, while our data support the importance of mtROS in driving tissue damage in neuroinflammation^6^, they expose microglia as a non-significant source.

### Reduction of mtROS in MdCs ameliorates neuroinflammation

We next explored whether reducing ROS production in MdCs could effectively ameliorate clinical symptoms and prevent tissue damage in EAE. To induce mCAT expression in MdCs, we crossed the *Ccr2^CreERT2^* strain to the mCAT^fl^ allele. EAE was induced in *Ccr2^CreERT^*^/+^ mCAT^fl/+^ mice and tamoxifen treatment was initiated at disease onset, as previously described (Fig. 4a). Immunoblot analysis confirmed overexpression of mCAT in MdCs (Extended Data Fig. 4a) along with a reduction in both cellular ROS (Fig. 4b) and mtROS (Fig. 4c). Importantly, this approach did not target microglia, and ROS levels in these cells remained unaltered (Fig. 4b-c).

**Figure 4.**
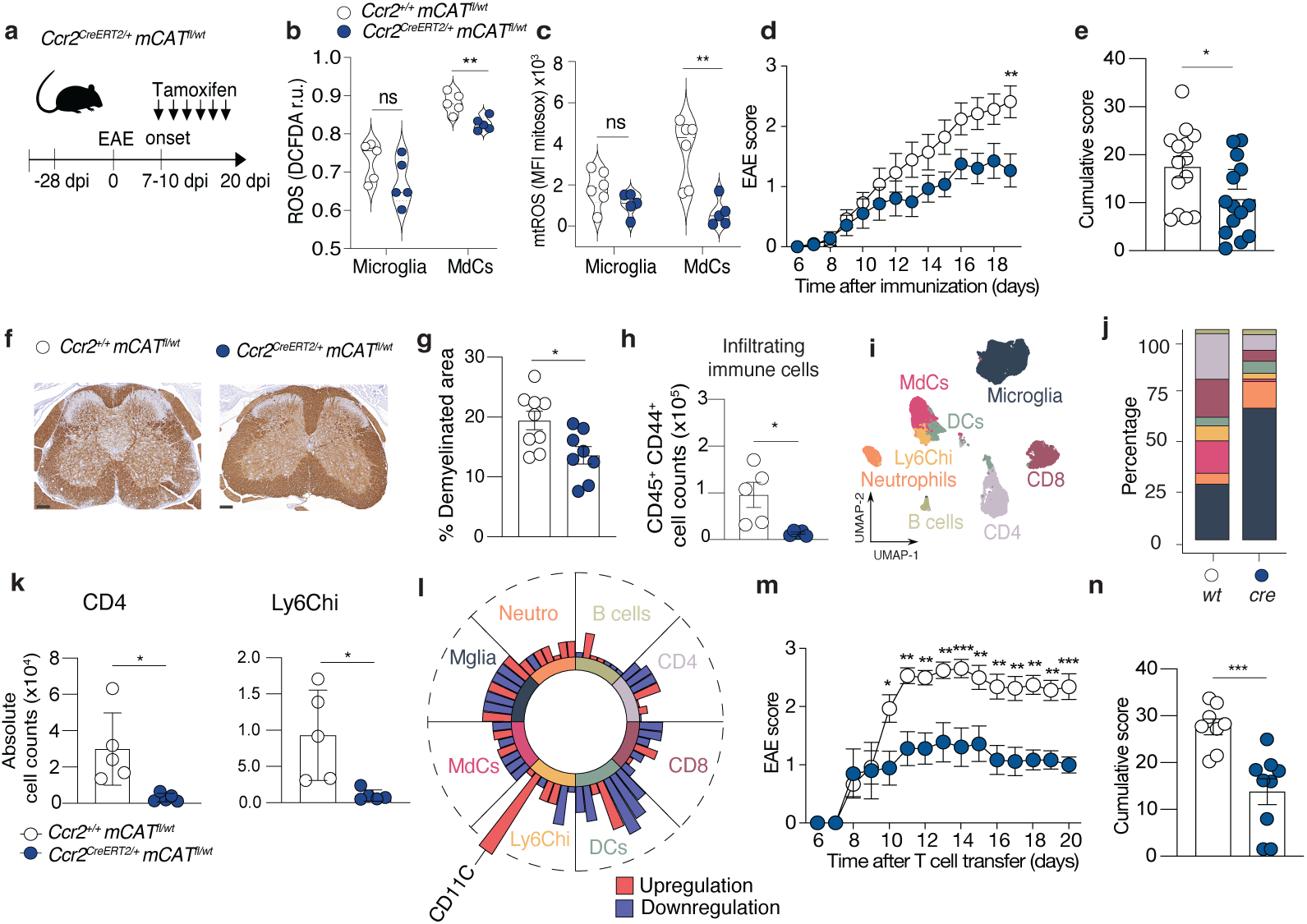
Quenching mtROS in MdCs allleviates neuroinflammation. **a**, Schematic representation of tamoxifen treatment to target bone marrow derived MdCs during EAE disease onset. **b**, ROS production measured by normalized DCFDA median expression in microglia and MdC clusters (generated in R) *Ccr2^+/+^*mCAT^fl/wt^ (n=5, m/f) and *Ccr2^CreERT2/+^* mCAT^fl/wt^ (n=5, m/f) mice. Pooled data from two experiments (unpaired two-tailed t-test; **P=0.0054 in MdCs). **c**, mtROS production measured as mitosox median expression in manually gated (FlowJo) microglia and MdCs from *Ccr2^+/+^*mCAT^fl/wt^ (n=6, m/f) and *Ccr2^CreERT2/+^* mCAT^fl/wt^ (n=5, m/f) mice. Pooled data from two experiments (unpaired two-tailed t-test, **P=0.003 in MdCs). **d**-**e**, Active EAE was induced by MOG_35-55_/CFA/PT immunization. Plots show mean clinical score over time (Two-way ANOVA, and Sidak’s post hoc test, day 19 **P=0.0078) (**d**) and cumulative clinical score (**e**) in *Ccr2^+/+^* mCAT^fl/wt^ (n=14, m/f) and *Ccr2^CreERT2/+^* mCAT^fl/wt^ (n=14, m/f) mice (unpaired two-tailed t-test; *P = 0.0339). Pooled data from two experiments. **f,** Representative MBP staining (left) and **g**, the percentage of myelin loss quantified in lumbar spinal cord sections in *Ccr2^+/+^* mCAT^fl/wt^ (n=9, m/f) and *Ccr2^CreERT2/+^* mCAT^fl/wt^ (n=8, m/f) mice (unpaired two-tailed t-test; **P = 0.0163). Data pooled from two independent experiments. **h,** Absolute counts of CD45+CD44+ CNS infiltrating immune cells (gated in FlowJo) at day 17 d.p.i. (n=5 per group, f). Unpaired two-tailed t-test (*P = 0.0164). **i,** UMAP from CNS leukocytes clustered by cell type and labelled based on surface marker expression. **j,** Bar graph of the percentage of the immune clusters found in in *Ccr2^+/+^* mCAT^fl/wt^ and *Ccr2^CreERT2/+^* mCAT^fl/wt^ (n=5 per group, f). **k**, Absolute counts of CNS infiltrating immune CD4 T cells (left) and Ly6Chi monocytes (right) (analysed in R) at day 17 d.p.i. (n=5 per group, f), (unpaired two-tailed t-test, *P=0.0386 in CD4 and *P=0.0403 in Ly6cHi). **l,** Radar plot showing changes in the marker expression of CNS immune cell clusters in *Ccr2^CreERT2/+^* mCAT^fl/wt^ relative to *Ccr2^+/+^* mCAT^fl/wt^ and (n=5 per group). Values correspond to −log10(p value) and were adjusted with the Benjamini-Hochberg correction. Bar color denotes the upregulation or downregulation of the respective marker. Marker order is consistent across subsets, left to right: CD38, MHCII, CD44, CX3CR1, F480, CD11C, and MerTK. Dashed line denotes −log10(0.05) cutoff for p value. **m-n** Passive EAE was induced by adoptive transfer of primed lymphocytes from actively immunized donor mice. Plots show (**m**) mean clinical score over time (two way ANOVA, and Sidak’s post hoc test, day 10 *P=0.0254, day 11 **P=0.0020, day 12 **P=0.0030, day 13 **P=0.0025, day 14 ***P=0.0006, day 15 **P=0.0076, day 16 **P=0.0019, day 17 **P=0.0020, day 18 **P=0.0013, day 19 **P=0.0039, day 20 ***P=0.0007) and (**n**) cumulative clinical score *Ccr2^+/+^* mCAT^fl/wt^ (n=8, m/f) and *Ccr2^CreERT2/+^* mCAT^fl/wt^ (n=9, m/f) mice (pooled data from two experiments, unpaired two-tailed t-test; ***P = 0.0009).

Intriguingly, blocking ROS production in MdCs lead to a significant reduction in EAE disease severity (Fig. 4d-e), demyelination (Fig. 4f-g) and immune cell infiltration (Fig. 4h) in *Ccr2^CreERT^*^/+^ mCAT^fl/+^ mice compared to tamoxifen treated littermate control mice. Spectral flow cytometry coupled with FlowSOM clustering revealed a change in the immune cell cluster composition (Fig. 4i-j), with a marked reduction in CD4^+^ T cells and Ly6C^hi^ monocyte cell counts (Fig. 4k). In contrast, no significant changes were observed in the MdC activation profile (Fig. 4l, Extended Data Fig. 4d).

To rule out any effects of mCAT overexpression on T cell priming, we induced passive EAE in *Ccr2^CreERT^*^/+^ mCAT^fl/+^ mice. This confirmed that mCAT overexpression in MdCs during the effector phase resulted in reduced disease severity (Fig. 4m-n). Collectively, these results demonstrate that invading MdCs, rather than microglia, are the primary ROS producers in neuroinflammation and play a critical role in driving tissue damage in CNS neuroinflammation.

## Discussion

CNS phagocytes are widely considered the primary mediators of tissue damage in both murine and human CNS neuroinflammation. A longstanding debate has centered on whether CNS-resident microglia or invading MdCs are to blame^16^. While many neurotoxic pathways have been proposed, none have been conclusively attributed to a specific phagocyte population. The higher abundance of microglia, their localization near lesions^7,14,29,17^, and their reactive phenotype during neuroinflammation^14,16,17,30^ have supported the idea that microglia are the main drivers of phagocyte-mediated tissue damage. However, the ability to distinguish microglia from MdCs and specifically target each population is beginning to challenge this view.

In this study, we characterized and selectively targeted oxidative stress in both microglia and MdCs. Our analysis of publicly available sequencing data revealed a conserved oxidative stress signature in MPs, specifically in bone marrow-derived MdCs during neuroinflammation. Functional experiments targeting ROS production suggest that invading MdCs, rather than embryonically derived microglia, are the primary contributors to ROS-mediated tissue damage.

The amelioration of EAE in mice lacking *Cybb,* the gene encoding NOX2, has been previously cited as evidence for microglia’s role in oxidative stress-mediated tissue damage^13,15^. Of note, NOX2 is involved in multiple cellular processes, including antigen processing^24,31^ and is expressed by various cell types implicated in neuroinflammation^24,31^. To clarify the role of NOX2 in microglia, we used a combination of conditional gene targeting and passive EAE to bypass its role in the priming of autoreactive T cells^31^. Surprisingly, we found that NOX2 deletion in MPs was insufficient to reduce ROS production or attenuate EAE severity. This aligns with recent transcriptional profiling of CNS phagocytes, which identified a core oxidative stress gene signature^3^ including several enzymes that drive ROS production. Thus, we conclude that NOX2 plays a redundant role in oxidative stress, and targeting it alone may not be enough to mitigate neuropathology in EAE. In contrast, general suppression of oxidative stress has shown neuroprotective effects in several models of neuroinflammation^3,6,10,11^. However, these studies, which relied on therapeutic compounds, did not pinpoint the specific cell types responsible for ROS-induced tissue damage.

Recent work has highlighted mitochondrial complex I (mtCI) activity as a key driver of reverse electron transport and pathogenic ROS production in CNS myeloid cells^6^. This activity was shown to sustain the DAM signature, and disrupting mtCI reduced EAE severity. While our findings support the functional importance of mitochondrial ROS (mtROS) in neuroinflammation, refined conditional gene targeting in mice challenges this interpretation by identifying MdCs, rather than microglia, as the primary cellular and molecular mediators of ROS-driven tissue damage. While ROS production may be crucial for MdC migration via the blood-brain barrier^33^, it can also cause direct oxidative damage in the CNS. Recent studies have shown that granulocyte-colony stimulating factor (GM-CSF)^29^, a key pathogenic T cell derived cytokine in EAE^32^ induces ROS production in MdCs. Thus, our findings provide functional evidence of how GM-CSF promotes MdCs pathogenicity in neuroinflammation.

In summary, by accurately identifying bone marrow-derived monocytic phagocytes as the main contributors to ROS-mediated tissue damage, our study resolves a longstanding controversy about the protagonists of oxidative stress in neuroinflammation. These findings shift the focus of neuroimmunology research and therapeutic intervention toward targeting MdCs, which may open new avenues for the treatment of CNS neuroinflammatory diseases.

## Methods

### Mice

All procedures were reviewed and approved by the Swiss Veterinary Office and performed according to institutional and federal guidelines. *Cx3cr1^CreERT2^* mice were originally provided by Steffen Jung and described in ^26^. *Ccr2*^CreERT2^ mice were described in ^23^. *Cybb*^fl^ mice were provided by Prof. Dr. Jan D. Lünemann, and originally described in ^34^. Mitocatalase (mCAT) mice were provided by A.S.S. and originally described in^27^. All transgenic mouse strains mice are bred in house on the C57BL/6 background and were maintained under optimized hygienic conditions (22°C; 45-65% humidity; 12 hours light cycle); standard laboratory diet and water were freely available.

### EAE induction

Active EAE was induced by injection of 200 μg of myelin oligodendrocyte glycoprotein epitope MOG_35-55_ (#RP10245, Genscript) emulsified in complete Freund’s adjuvant (CFA) (#231131, BD). Mice received two subcutaneous (s.c.) injections into the flank and intraperitoneal injection (i.p.) of 200 ng pertussis toxin (PT) (List Biological Laboratories, #179A) the same day and two days after.

For passive EAE, splenocytes and lymph node cells were isolated from MOG_35-55_/CFA immunized donor mice (day 11 p.i.) that did not receive PT. Cells were cultured in medium supplemented with IL-23 (1887-ML-010/CF, R&D, 20 ng/ml), anti-IFNγ (505713, BioLegend, 10ug/ml) and MOG_35-55_ peptide (GenScript, 10ug/ml). After 3-4 days in culture 3.5-5x10^6^ cells were injected i.p..

EAE clinical scores were defined as follows: no detectable signs of EAE: 0; tail limp at distal end: 0.5; entirely limp tail: 1; limp tail and hind limb weakness (occasional grid test positive): 1.5; unilateral partial hind limb paralysis (one leg constantly falls through the grid: 2; bilateral partial hind limb paralysis: 2.5; complete bilateral hind limb paralysis: 3; complete bilateral hind limb paralysis and partial forelimb paralysis: 3.5; complete paralysis of fore and hind limbs (moribund): 4. In case the clinical symptoms couldn’t be assigned to one or the other score, we used “in between” scoring intervals of 0.25 which allowed a more accurate description of the clinical symptoms. Withdrawal criteria included: clinical score ≥ 3 for more than 7 days, clinical score ≥ 3.5 for more than 3 days, weight loss > 25%, as well as pain symptoms.

### Tamoxifen treatment

Tamoxifen (Sigma) was dissolved in ethanol and corn oil to 25 mg/ml and administered in 200 μl doses via oral gavage (5 mg per dose). Tamoxifen schemes are described for each strain.

### Cell isolation from the adult mouse CNS

Mice were sacrificed through CO_2_ asphyxiation and transcardially perfused with ice-cold PBS. For leukocyte isolation, whole brain and spinal cord was harvested, cut into small pieces and digested with 0.4 mg/ml Collagenase IV (#9001-12-1, Sigma-Aldrich) and 0.2 mg/ml deoxyribonuclease I (DNase I) (#E1010, Luzerna) in HBSS (with Ca^2+^ and Mg^2+^) (#14025-050, Gibco) for 40 min at 37°C or processed in the gentleMACS™ Octo Dissociator with Heaters program 37C_ABDK_01. The digested tissue was then mechanically dissociated using a 19-gauge needle and filtered through a 100 μm cell strainer (800100, Bioswisstec). CNS single-cell suspensions were further enriched by 30 % Percoll^®^ (P4937, GE Healthcare) gradient centrifugation (2750 rpm, 30 min, at 4°C, no brakes).

### FC surface staining

Prior to surface labelling, cells were incubated with purified anti-mouse CD16/32 (clone 93, BioLegend) for 10 min on ice to prevent non-specific binding of primary antibodies. Single-cell suspensions were then directly incubated with the primary surface antibody cocktail for 20 min at 4°C. Cells were washed with FACS buffer and incubated in the secondary surface antibody cocktail (fluorochrome-conjugated streptavidin) for 20 min at 4°C.

Anti-mouse fluorochrome-conjugated monoclonal antibodies used in this study were CD274 (78-5982-62, eBioscience, clone MIH5, SuperBright780, dilution 1:200), CD38 (102746, BioLegend, clone 90, APC-Fire 810, dilution 1:400), I-A/I-E (107620, BioLegend, clone M5/114.15.2, PB, dilution 1:400), Siglec-F (562757, BD, clone E50-2440, PE-CF594, dilution 1:200), CD45 (564279, BD Biosciences, clone 30-F11, BUV395, dilution 1:200), Ly6G (565707, BD Biosciences, clone 1A8, BUV563, dilution 1:100), CD19 (565076, BD Biosciences, clone 1D3, BUV661, dilution 1:200), CD44 (612799, BD Biosciences, clone IM7, BUV737, dilution 1:200), CD64 (139309, BioLegend, clone X54-5/7.1, BV421, dilution 1:100), F4/80 (123112, BioLegend, clone BM8, dilution 1:500), CX3CR1 (149027, BioLegend, clone SA011F11, BV605, dilution 1:400), CD11b (101239, BioLegend, clone M1/70, BV510 dilution 1:400), Ly-6C (128037, BioLegend, clone HK1.4, BV711, dilution 1:400), MerTK (PECy7,), CD11c (35-0114-82, eBioscience, clone N418, PE-Cy5.5, dilution 1:800), CD88 (135811, BioLegend, clone 20/70, Biotin, dilution 1:200), CD3 (100236, BioLegend, clone 17A2, APC, dilution 1:200), CD26 (137804, BioLegend, clone H194-112, PE, dilution 1:100), CD49d (741925, BD Biosciences, clone R1-2, BUV 805, dilution 1:100).

### DCFDA (ROS) staining

After FC surface staining, CNS single-cell suspensions were stained with 5 mM of DCFDA (Abcam, Ab113851) for 30 min at 37°C or 4 °C. Next, cells were washed with PBS 1X and inmediately acquired.

### Mitosox (mROS) staining

After FC surface staining, CNS single-cell suspensions were stained with 5 μM of mitoSOX mitochondrial indicator (Thermo Fisher Scientific, M36008) for 20 min at 37 °C. Next, cells were washed three times with pre-warmed HBSS and immediately acquired.

### FC data acquisition and analysis

Samples were acquired on the 5L Aurora spectral analyser (Cytek Biosciences) and data analyzed using the FlowJo software (version 10.8.0, Tree Star Inc.).

### Fluorescence-activated cell sorting

Cell sorting of live CD11b^+^CD45^+^CX3CR1^+^CD44^-^ microglia and CD11b^+^CD45^+^ CX3CR1^+^CD44^+^Ly6C^+^Ly6G^-^ MdCs was performed using a BD FACS Aria III (FACS DIVA Software v9). Anti-mouse fluorochrome-conjugated monoclonal antibodies used in this study were: CD45 (103126, BioLegend, clone 30-F11, Pacific Blue, dilution 1:600), CD11b (101263, BioLegend, clone M1/70, BV510, dilution 1:400), CD44 (103049, BioLegend, clone IM7, BV650, dilution 1:400), CX3CR1 (149027, BioLegend, clone SA011F11, BV605, dilution 1:400), Ly6C (128037, BioLegend, clone HK1.4, BV711, dilution 1:400), Ly6G (127606, BioLegend, clone 1A8, FITC, dilution 1:400).

### Quantitative RT–PCR

Total RNA was isolated from sorted microglia and MdCs using the QuickRNA Microprep Kit (R1051, Zymo Research) according to the manufacturer’s instructions. Complementary DNA (cDNA) was synthesized using the M-MLV Reverse Transcriptase (28025013, Invitrogen) and qRT–PCR was performed on a CFX384 Touch Real-Time PCR Detection System (Bio-Rad) using SYBR Green (Bio-Rad). Gene expression was calculated as 2−ΔCt relative to Pol2 as the endogenous control. Relative gene expression was then calculated by dividing cre gene expression value by the average of wt gene expression value.

### Primers for Cybb

5′-ACTCCTTGGGTCAGCACTGG-3′; (forward)

5′-GTTCCTGTCCAGTTGTCTTCG-3′; (reverse)

### Primers for Pol2

5′-CTG GTC CTT CGA ATC CGC ATC-3′; (forward)

5′-GCT CGA TAC CCT GCA GGG TCA-3′; (reverse)

### Immunoblot analysis

Sorted microglia and MdCs were lysed with Laemli buffer and boiled for 5 min at 95°C. Cell lysates were separated in 10% SDS-PAGE gels and transferred onto PVDF membranes (Whatman). The membranes were blocked with PBS-Milk 5% for 30 min at RT before staining with primary antibody against HA-tag (C29F4, Cell signaling, 1:500) and Actin (ab49900, Abcam, 1:500) overnight at 4°C and secondary antibody for 1h at RT. Finally, membranes were developed with ECL detection kit (WesternBright Sirius femtogram HRP Substrate Witec AG7002325).

### Histopathological and Immunohistochemical Quantitative Analysis of Spinal Cords

Mice were euthanized through CO2 inhalation and transcardially perfused with ice-cold PBS. Spinal columns were carefully dissected and fixed in 4% PFA (11762.00500, Morphisto) or 10% neutral formalin for 24 hours at 4°C, rinsed in PBS followed by decalcification: in 0.5M EDTA solution (A3145.0500, Axonlab) at pH 8, at 4°C for five consecutive days or decalcified in RDFTM (Biosystems, article no. 81-0835-00; 10-20% formic acid) for 48 h decalcified in RDFTM (Biosystems, 81-0835-00; 10-20% formic acid) in rotation. Tissue dehydration was performed in graded alcohols. Transversal lumbar sections of the spinal cord were dissected and embedded in paraffin. 3µm-thick tissue sections were cut on a microtome (Microm HM 360, Thermo Fisher) and transferred to mounted Superfrost^TM^ Plus microscope slides (Carl Roth, Germany). Serial sections were prepared for hematoxylin and eosin (HE) staining as well as immunohistochemistry (IHC). For IHC using the horseradish peroxidase (HRP) method, sections of the lumbar spinal cord segments were stained with a primary antibody targeting myelin basic protein (MBP). Briefly, deparaffination was followed by 1-hour incubation in room temperature with the primary antibody (monoclonal rat, MAB386; Merck Millipore, Darmstadt, Germany) diluted 1:800 in dilution buffer (Agilent Dako, Glastrup, Denmark). Subsequently, an incubation with the appropriate secondary antibody/detection system (rabbit-anti-rat, article no. BA 4000, Vector Laboratories, Newark, USA; EnVision+ rabbit, Agilent Dako, Glastrup, Denmark) in an autostainer (Agilent Dako, Glastrup, Denmark). Sections were subsequently counterstained with hematoxylin.

IHC and HE images were acquired using NanoZoomer 2.0-HT; Hamamatsu, Hamamatsu City, Japan or Zeiss Axio Scan. Z1 Slidescanner (Zen 2 software, blue edition) with 10x or 20x objectives. The scanned slides of each animal were quantitatively analyzed using the Visiopharm 2022.01.3.12053 software (Visiopharm, Hoersholm, Denmark).

To quantify demyelinated area, regions of interest (ROIs) corresponding to the white matter coverage of the spinal cord were drawn. Within these ROIs, the MBP positive area (MBP+%) was calculated. Demyelinated area was defined as total white matter area of the spinal cord circumference-MBP+%. We then report the average percentage demyelination of each mouse.

### Analysis of publicly available datasets

Publicly available single cell (sc) and single nuclei (sn) RNA-seq datasets were analyzed using Seurat 5.1.0 in R 4.4.0. Cells were subsetted based on immune-related annotations from the original datasets. Specifically, we kept/were kept cells annotated as “immune” and “lymphocytes” from Absinta *et al* (GEO: GSE180759); “macrophages and microglia”, “macrophages” and “immune cells” from Jäkel *et al* (GEO: GSE118257); “microglia”, “phagocytes”, “T cells” and “B cells” from Schirmer *et al* (SRA: PRJNA544731); “microglia”, “T cells” and “B cells” from Macnair *et al* (Zenodo: 10.5281/zenodo.8338963); and “myeloid cells” and “T cells” from Kiss *et al* (GEO: GSE227781). First, principal component analysis (PCA) was performed on the 2000 most variable features. To correct for batch effect, we then integrated the datasets using the Harmony method7. Subsequently, Louvain community clustering was applied to a 20-nearest neighbor graph generated using the harmony embedding/reduction Clusters were annotated as follows: BAMs = 14; MPs = 15, 3, 0, 5, 6, 8, 19, 20, 2, 9, 4, 10, 18,1, 13; Other = 21, 12, 17, 11; Lymphocytes = 7, 16. A uniform manifold approximation and projection (UMAP) was created with the first 30 components of the Harmony embedding.

Samples across the different datasets belonging to non-MS patients were grouped as “control”. White matter and grey matter were categorized separately. White matter was subdivided into active, chronic active, chronic inactive, edge, and remyelinating lesions, as well as normal-appearing regions. Lesions originally annotated as “active lesion”, “lesion core”, and “plaque” were grouped under the active lesion category. “Acute chronic active” and “chronic active” lesions were also combined. The lesion edge category included “chronic active lesion edge” and “chronic inactive lesion edge. “ Normal-appearing white matter encompassed “normal appearing”, “MS white matter”, and “periplaque white matter.” For grey matter, “normal appearing grey matter” and “MS grey matter” were combined, while grey matter lesions were not further subdivided in the original datasets.

Publicly available scRNA-seq mouse datasets from Mendiola *et al* (GEO:GSE146295)^3^, Jordão *et al* (GEO: GSE118948)^22^, and Peruzzotti-Jametti *et al* (GEO:GSE248175)^6^ were each individually re-clustered using the Louvain algorithm and manually re-annotated with the same homeostatic microglia (hMicroglia) (*P2ry12, Tmem119, Sall1*), DAM (*C1qa, C1qc, Cst7, Ccl12 and Ly86*) and MdC (*Ccr2, Ly6c2, Cd14, H2-Ab1, Clec4e and Arg1*) marker genes to ensure consistency across all datasets.

The AddModuleScore function in Seurat was used to compute cell-type-specific gene set activity scores, or module scores. Module scores for reactive oxygen species (ROS) were calculated using the GO:0072593 ROS metabolic process gene set (as used in Mendiola *et al*^3^), with the mouse-specific gene set used for mouse datasets and human-specific gene set used for human datasets. For mitochondria complex I, module scores were computed using the same genes as described in Peruzzotti-Jametti *et al* ^6^.

### High dimensional analysis of FC data

Compensated and cleaned data was exported from FlowJo to R studio version 4.0.1. After transformation and percentile normalization, UMAPs were generated using the package ‘umap’ version 0.2.7.0. Data were clustered using FlowSOM version 2.6.0 and “ConsensusClusterPlus” package subsequently. Metaclusters were merged and annotated based on the median expression profile and their localization on the UMAP. Heatmaps were generated using the pheatmap’ package version 1.0.12.

### Statistical Analysis

To assess data normal distribution, the Shapiro–Wilk test or Kolmogorov–Smirnov test were used and the quantile–quantile plots were examined. Group means were compared with two-tailed, unpaired t-tests or Mann–Whitney tests depending on data normal distribution. Multiparametric analysis was performed with two-way analysis of variance (ANOVA), followed by Sidak’s multiple comparison tests. P values below 0.05 were considered as significant and are indicated by asterisks (*P < 0.05, **P < 0.01, ***P < 0.001) and numerical values are reported on the respective figure legends. *N* represents the number of biologically independent animals. Data are presented as mean ± standard error of the mean (s.e.m) or as mean± standard deviation (s.d) as reported on the respective figure legends. Statistical analysis was carried out using GraphPad Prism 9 (GraphPad Software, Inc.) or in R Corrections for multiple hypothesis testing were done using the Benjamini-Hochberg method. Statistical methods are indicated in the figure legends.

### Data and code availability

Data and code will be available upon request.

## Acknowledgments

This work has received funding from the Swiss National Science Foundation (Ambizione PZ00P3_193330 to S.M. and Eccellenza PCEFP3_187000 to A.S.S.), the Research Talent Development Fund zur Förderung des akademischen Nachwuchses (FAN; to S.M.), Dr. Wilhelm Hurka Foundation (to B.B., D.D.F. and S.M.), the Swiss Multiple Sclerosis Society (to S.M. and D.D.F.), the Hartmann-Müller Foundation (to S.M.), the Novartis Foundation for Medical-Biological Research to S.M. and the UZH Candoc grant (FK-22-040 to J.V-V.).

## Author contributions

J.V-V., S.M. and D.D.F. designed and performed all mouse experiments. J.V-V. analyzed and interpreted all data and prepared all Figures. P.C. and J.V-V. performed analysis of public snRNAseq and scRNAseq data. M.P-F. J.K., E.M., V.B., D.B., L.O., K. W. S., S.C-P., U.D helped with experiments. F.I. and C.U. helped with computational data analysis and provided code. F.S., F.P. and and M.P-F. performed IHC and immunoblot stainings and quantitative immunohistochemical analysis. S.M., J.V-V., B.B., U.D., D.D.F., C.M., A.S.S. provided scientific and intellectual input. J.V-V., P.C. and S.M. wrote and D.D.F, F.I., C.M and B.B. edited the manuscript. S.M. supervised and aquired funding for the study. All authors approved the paper.

---

**Extended Data Fig. 1.**
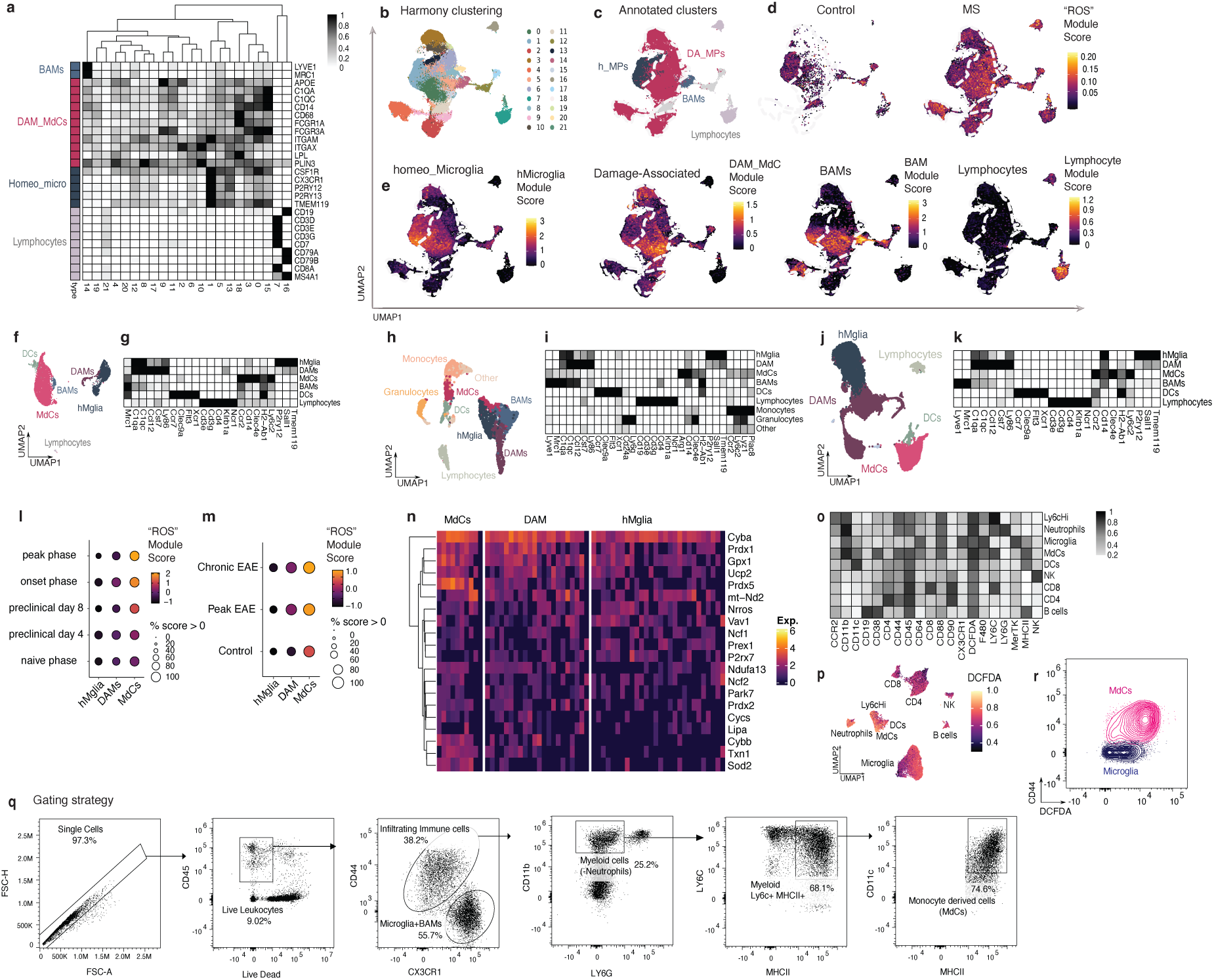
**a** Heatmap of markers expressed by the different clusters and their annotations. **b,** Harmony-integrated Louvain clusters identified in the dataset. **c,** UMAP of the annotated immune clusters. **d,** UMAPs separated by control and MS with overlay of the “ROS” module score. **e,** UMAP overlay of the genes used to annotate Homeostatic Mononuclear Phagocytes (MPs), Damage-associated (DA) MPs, and Monocyte-derived MPs. **f,** UMAP and **g,** Heatmap from the Mendiola *et al.*^3^ dataset. **h**, UMAP and **i,** Heatmap from the Jordao *et al* ^22^ dataset. **j**, UMAP and **k,** Heatmap of cluster expression for the annotated clusters in the Peruzzotti-Jametti *et al* ^6^ dataset. **l,** Dot plot of the “ROS” module score at different stages of EAE from the Jordao *et al* ^22^ dataset. **m,** Dot plot of the “ROS” module score during control, peak, and chronic EAE stages in the Peruzzotti-Jametti *et al* ^6^ dataset. **n**, Clustered heatmap of the 20 most expressed ROS-associated genes in MPs found in the Peruzzotti-Jametti *et al* ^6^ dataset. **o**, Heatmap of marker expression from annotated immune clusters during peak EAE. **p,** Overlayed ROS (DCFDA) expression. **q,** Dot plots depict the gating strategy used to calculate the infiltrated immune cell counts, Live CD45+CD44+, DCFDA and Mitosox expression in Microglia (Microglia+BAMs) and Monocyte derived cells (MdCs). **r,** Biaxal contour plot of CD44 (y-axis) and DCFDA (x-axis) expression in MdCs and microglia. For panels **l** and **m**, circle size depicts the percentage of expression detected per cluster, and color indicates the expression level as the average module score or average expression.

**Extended Data Fig. 2.**
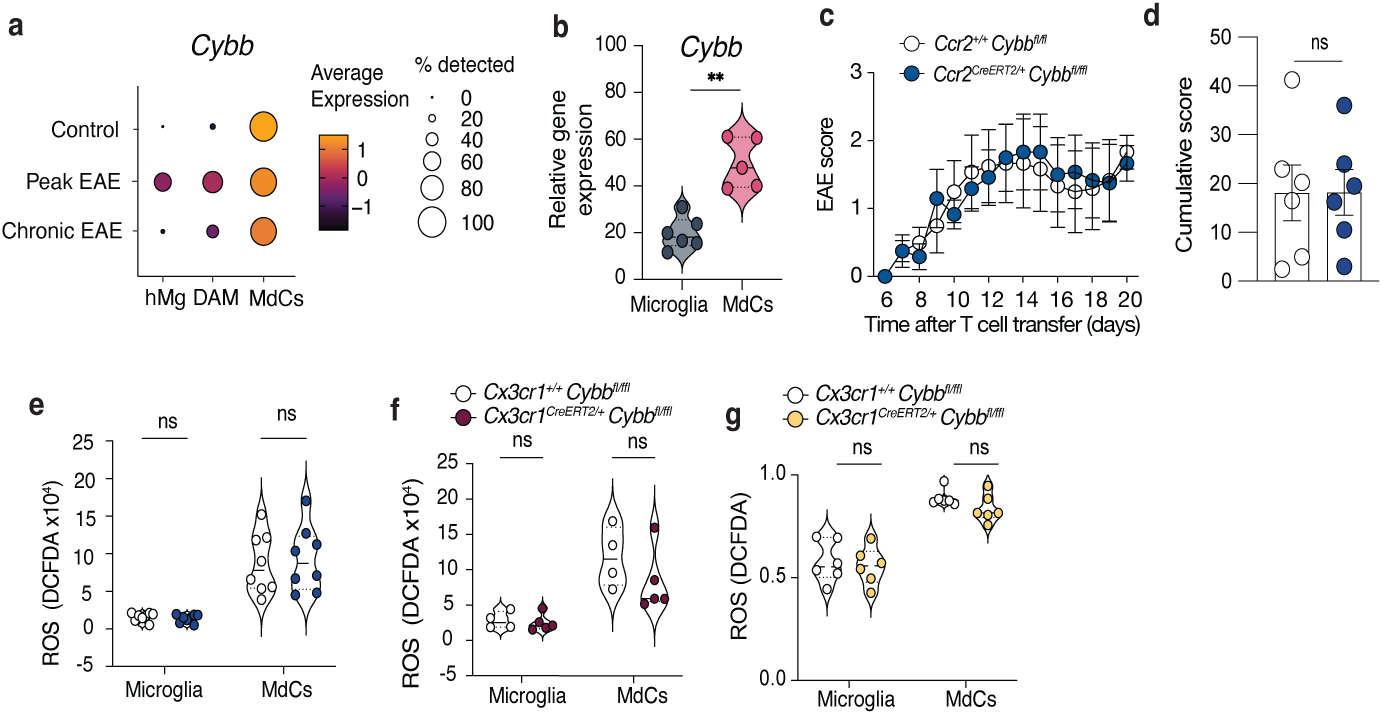
**a,** Dot plot illustrating *Cybb* expression in homeostatic microglia (hMicro), damage-associated microglia (DAMs), and monocyte-derived cell clusters (MdCs) from Peruzzotti-Jametti *et al*^6^ during control (non-immunized), peak and chronic phases of EAE. Circle size is depicting % expression detected per cluster and color is depicting expression level as Average Module Score or Average Expression. **b,** Relative gene expression *Cybb* measured by qPCR of FC-sorted Microglia (n=6) and MdCs (n=5) during peak EAE. (Mann–Whitney test; **P = 0.0022). **c,d,** Passive EAE was induced by adoptive transfer of primed lymphocytes from actively immunized donor mice. Plots depict mean clinical score (**c**) overtime (two-way ANOVA, and Sidak’s post hoc test) and (**d**) cumulative clinical score *Ccr2^+/+^ Cybb^fl^*^/fl^ (n=5, f) and *Ccr2 ^CreERT2/+^ Cybb^fl^*^/fl^ (n=6, f) mice (unpaired two-tailed t-test). **e,** ROS production measured by DCFDA median expression in manually gated (FlowJo) microglia and MdC from *Ccr2^+/+^ Cybb^fl^*^/fl^ (n=7, m/f) and *Ccr2 ^CreERT2/+^ Cybb^fl^*^/fl^ (n=8, m/f) mice (unpaired two-tailed t-test). **f,** ROS production measured by DCFDA median expression in manually gated (FlowJo) microglia and MdC from *Cx3cr1^+/+^ Cybb^fl^*^/fl^ (n=4, m/f) and *Cx3cr1 ^CreERT2/+^ Cybb^fl^*^/fl^ (n=5, m/f) mice. Tamoxifen regimen corresponds to early tamoxifen treatment (Fig. 2f) (unpaired two-tailed t-test). **g,** ROS production measured by DCFDA median expression in microglia and MdC clusters (generated in R) in *Cx3cr1^+/+^ Cybb^fl^*^/fl^ (n=6, m/f) and *Cx3cr1 ^CreERT2/+^ Cybb^fl^*^/fl^ (n=6, m/f) mice (unpaired two-tailed t-test).

**Extended Data Figure 3.**
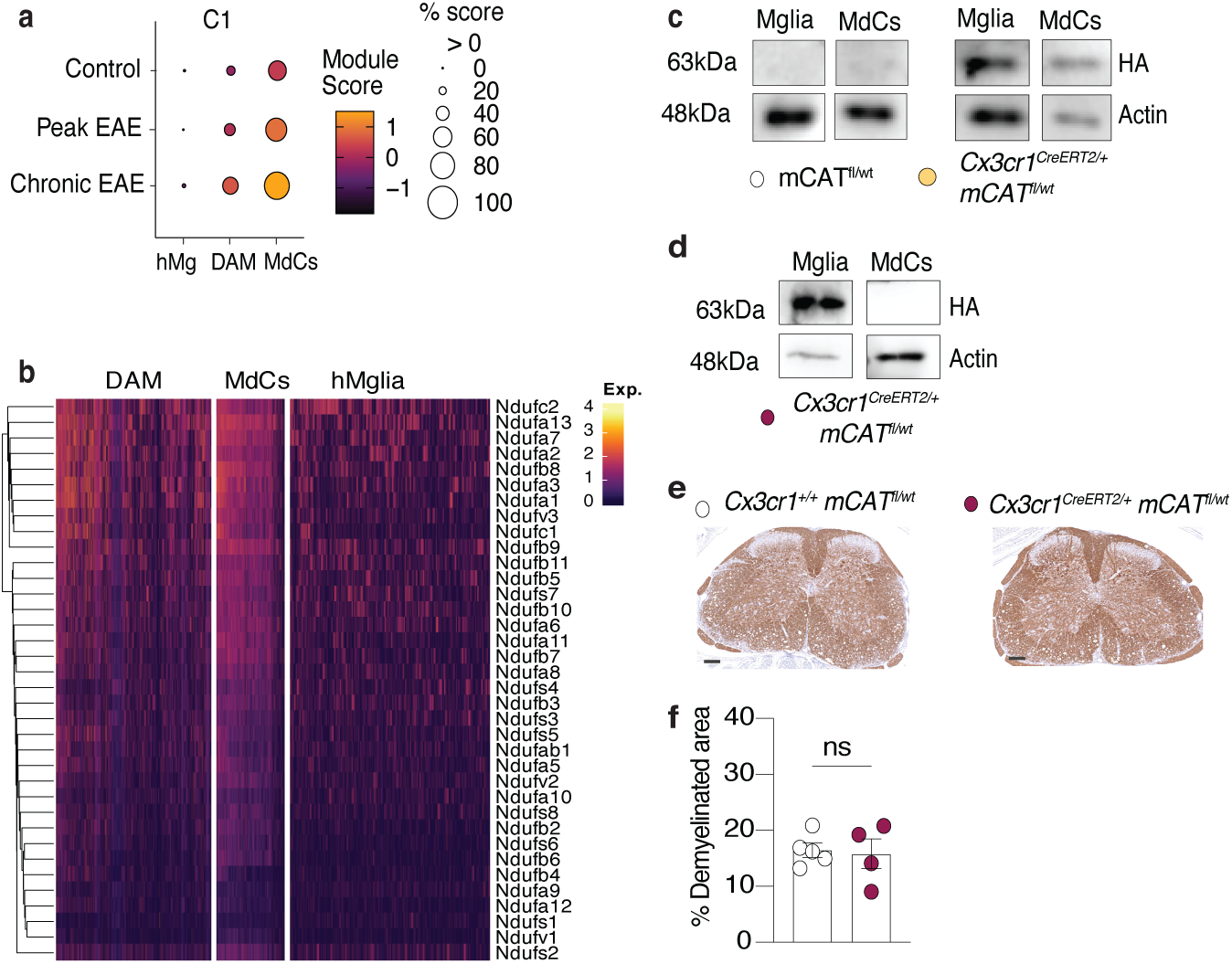
**a**, DotPlots illustrating mitochondria complex I module score in CNS microglia and MdCs in the Peruzzotti-Jametti *et al*. ^6^ dataset in peak and chronic EAE compared to control CNS. Circle size is depicting % expression detected per cluster and color is depicting expression level as Average Module Score. **b**, Clustered heatmap of the expression of the mitochondria complex I genes in MPs found in the Peruzzotti-Jametti *et al*. ^6^ dataset. **c**, Immunoblot showing HA-tag and actin bands, in FACs sorted microglia and MdCs from *Cx3cr1^+/+^*mCAT^fl/wt^ and *Cx3cr1^CreERT2/+^* mCAT^fl/wt^ mice (pooled cells from n=3 mice, m/f). **d**, Immunoblot showing HA-tag and actin bands, in FACs sorted microglia and MdCs early treatment tamoxifen regimen (Fig. 3g) *Cx3cr1^CreERT2/+^* mCAT^fl/wt^ mice (pooled cells from n=2 mice, m/f). **e-f**, Representative MBP staining (left) and the percentage of myelin loss quantified in lumbar spinal cord sections in early treatment tamoxifen regimen (Fig. 3g) *Cx3cr1^+/+^* mCAT^fl/wt^ (n=5, f) and *Cx3cr1^CreERT2/+^* mCAT^fl/wt^ (n=4, f) mice (right). Unpaired two-tailed t-test.

**Extended Data Figure 4.**
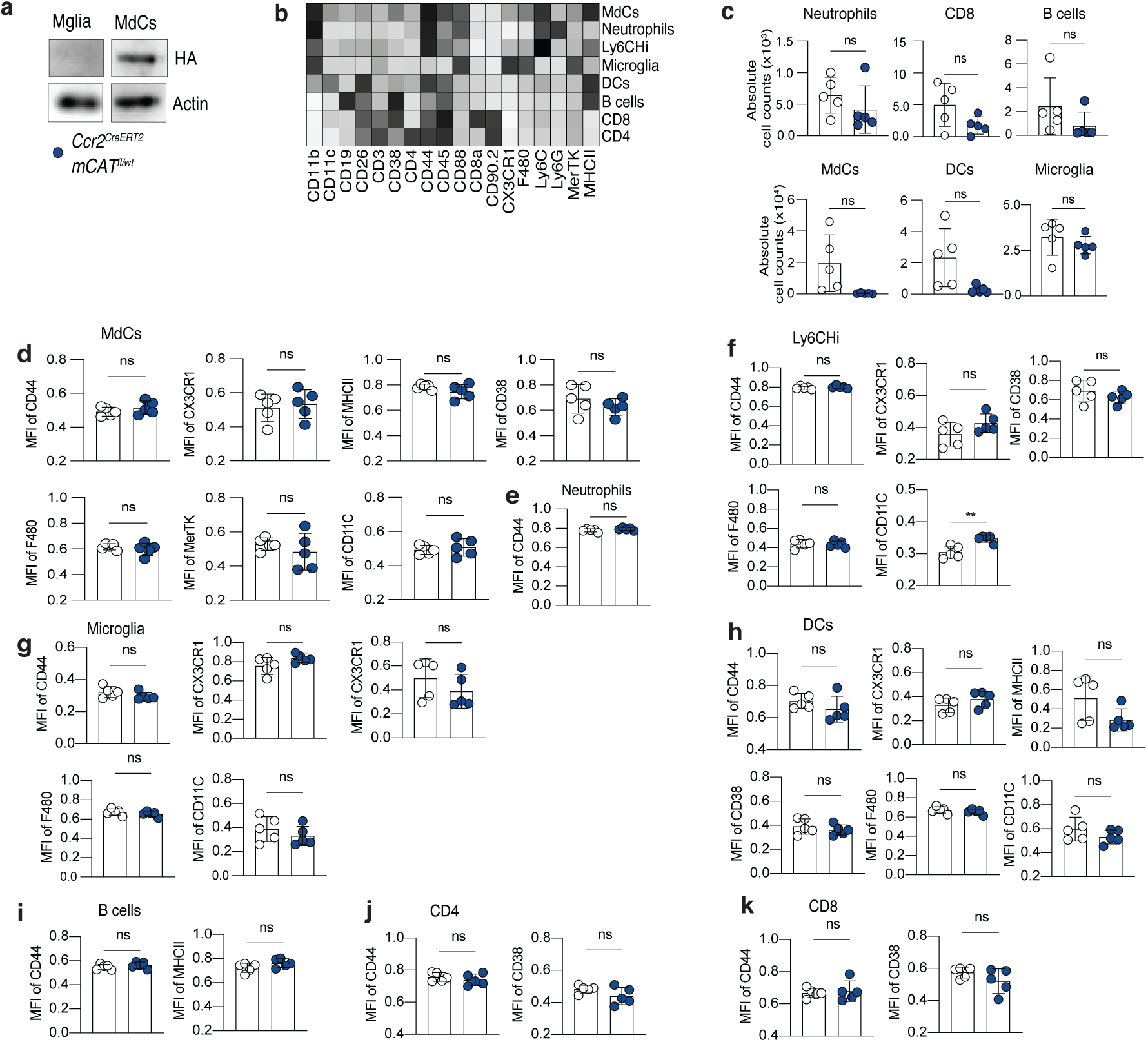
**a**, Immunoblot showing HA-tag and Actin bands, in FACS sorted microglia and MdCs from *Ccr2^CreERT2/+^* mCAT^fl/wt^ (pooled cells from n=3 mice). **b**, Heatmap showing the median marker expression across the identified main clusters, UMAP is shown in Fig. 4h. **c,** Absolute counts of CNS infiltrating Neutrophils, CD8 T cells, B cells, Monocyte derived cells (MdCs), Dendritic cells (DCs) and CNS resident immune Microglia (R generated). **d-k** Median fluorescent intensity (MFI) of activation markers (R generated) in (**d**) MdCs, (**e**) Neutrophils, (**f**) Ly6cHi Monocytes, (**g**) Microglia, (**h**) Dendritic cells, (**i**) B cells, (**j**) CD4 T cells and (**k**) CD8 T cells. (For MFI of CD11C in Ly6cHi Monocytes, unpaired two-tailed t-test. **P=0.0066). Mice analyzed at 17 d.p.i (n=5 per group, f), (unpaired two-tailed t-test).

